# BRAINformat: A Data Standardization Framework for Neuroscience Data

**DOI:** 10.1101/024521

**Authors:** Oliver Rübel, Prabhat, Peter Denes, David Conant, Edward Chang, Kristofer Bouchard

## Abstract

Neuroscience is entering the era of ‘extreme data’ with little experience and few plans for the associated volume, velocity, variety, and veracity challenges. This is a serious impediment for both the sharing of data across labs, as well as the utilization of modern and high-performance computing capabilities to enable data driven discovery. Here, we introduce BRAINformat, a novel file format and model for management and storage of neuroscience data. The BRAINformat library defines application-independent design concepts and modules that together create a general framework for standardization of scientific data.

We describe the formal specification of scientific data standards, which facilitates sharing and verification of data and formats. We introduce the concept of *Managed Objects*, enabling semantic components of data formats to be specified as self-contained units, supporting modular and reusable design of data format components and file storage. The BRAINformat is built off of HDF5, enabling portable, scalable, and self-describing data storage. We introduce the novel concept of *Relationship Attributes* for modeling and use of semantic relationships between data objects, and discuss the annotation of data using dedicated data annotation modules provided by the BRAINformat library. Based on these concepts we implement dedicated, application-oriented modules and design a data standard for neuroscience data. The BRAINformat software library is open source, easy-to-use, and provides detailed user and developer documentation and is freely available at: https://bitbucket.org/oruebel/brainformat.

## 1 INTRODUCTION

Neuroscience research is facing an increasingly challenging ‘big data’ problem due to the growing complexity of experiments and the volume/variety of data being collected from many acquisition modalities. Neuroscientists are routinely collecting data in a broad range of data formats that are often highly domain specific, ad-hoc and/or designed for efficiency with respect to very specific tools and data types. Even for single experiments, scientists are interacting with often tens of different formats—one for each recording device and/or analysis—while many data standards are not well-described or are only accessible via proprietary software. Navigating this quagmire of formats hinders efficient data analysis, data sharing, and collaboration and can lead to errors and misinterpretation of data. File formats and data standards that can represent complex neuroscience data and make the data easily accessible play a key role in enabling scientific discovery, development of reusable tools for data analytics, and progress towards fostering collaboration in the neuroscience community.

The requirements towards a data format standard for neuroscience are highly complex and go far beyond the needs of traditional, data modality-specific formats (e.g. image, audio, or video formats). A neuroscience data format needs to support the management and organization of complex collections of data from many modalities and sources, e.g., neurological recordings, audio and video recordings, eye-tracking, motion tracking, task contingencies, external stimuli, derived analytic results, and many others. To enable data interpretation and analysis, the format needs to also support storage of complex metadata, e.g., descriptions of recording devices, experiments, subjects etc..

Advanced neurosciences analytics furthermore rely on complex data access patterns driven by data semantics. For example, to study human brain activity underlying speech, scientists need to be able to efficiently annotate and extract data using complex combinations of annotations. Annotating data in itself, however, is a highly complex task that requires the coordinated access to related data sources. For example, a scientists may use audio or video recordings to identify particular events of interest and in turn needs to locate the corresponding data in a neural recording dataset to annotate it. Therefore, it is crucial that neuroscience formats support annotation of data as well as the specification and use of relationships between data objects.

In addition to these more application-specific needs, a usable, sustainable, and extensible data format also needs to satisfy a broad range of general, advanced file format and API requirements — e.g, the format should be self-describing, easy-to-use, efficient, portable, scalable, verifiable, easy to share and should support self-contained and modular storage of large data. Meeting all these complex needs is a daunting challenge. Arguably, the focus of a neuroscience-oriented data standard should be on addressing the application-centric needs of organizing scientific data and metadata, rather than on reinventing file storage and format methods. For the development of BRAINformat we have utilized HDF5 as the basic storage format as it already satisfies a broad range of the more basic format requirements—HDF5 is self-describing, portable, extensible, widely supported by programming languages and analysis tools, and is optimized for storage and I/O of large-scale scientific data.

In this manuscript we introduce the BRAINformat, a novel data format standardization framework and API for scientific data, developed at the Lawrence Berkeley National Labs in collaboration with neuroscientists at UCB and UCSF. BRAINformat supports the formal specification and verification of scientific data formats and supports the organization of data in a modular, extensible, and reusable fashion via the concept of *managed objects* (Sec. 3.1). To enable the modeling and use of complex relationships between data objects, we introduce the novel concept or *relationship attributes*. Relationship attributes support the specification of structural and semantic links between data, enabling users and developers to formally document and utilize relationships in a well-structured and programmatic fashion (Sec. 3.2). We demonstrate the use of chains of relationships to model complex relationships between multi-dimensional arrays based on data registration via the concept of advanced *index map relationships* (Sec. 3.2.4). The BRAINformat library and format also provides advanced support for definition, storage, and management of complex collections of data annotations (Sec. 3.3). We demonstrate the application of our framework to design a novel data standard for neuroscience data and its application to the storage and management of electrocorticography data collected during speech production (Sec. 4).

## 2 BACKGROUND AND RELATED WORK

The scientific community utilizes a broad range of data formats, which typically focus on different levels of the data organization and storage problem. Basic formats explicitly specify how data is laid out and formatted in binary or text data files (e.g., CSV, BOF, etc). While such basic formats are common in practice, they generally suffer from a lack of portability, scalability and a rigorous specification. For text-based files, languages and formats, such as the Extensible Markup Language (XML) [2] or the JavaScript Object Notation (JSON) [1], have become popular means to standardize documents for interchange of data. XML, JSON and other text-based data standards (in combination with character-encoding schema, e.g, ASCII or Unicode) play a critical role in practice in the exchange of usually relatively small, structured documents but are impractical for storage and exchange of large scientific data arrays due to the large overheads and cost they entail for storage, transfer, and I/O.

For storage of large-scale scientific data, HDF5 [14] and NetCDF [10] among others, have gained wide popularity. HDF5 is a data model, library, and file format for storing and managing large and complex data. HDF5 defines a set of core data object primitives, specifically: i) *Groups* which are similar to folders on a file system, used to group data objects, ii) *Datasets* which define n-dimensional arrays of arbitrary shape and data type, and iii) *Attributes* which are small meta data objects describing the nature and/or intended usage of groups or datasets. These data object primitives in combination provide the foundation for the organization and storage of highly complex data. HDF5 is portable, scalable, self-describing, and extensible and is optimized for storage and I/O of large-scale data. HDF5 is widely supported across programming languages and systems—e.g. R, Matlab, Python, C, Fortran, VisIt, ParaView etc.—and the HDF5 technology suite includes tools and applications for managing, manipulating, viewing, and analyzing data in the HDF5 format. HDF5 has been adopted as a base format across a broad range of application sciences, ranging from physics to bio-sciences and beyond^1^. Self-describing formats like HDF5 address the critical need for standardized storage and exchange of complex and large scientific data.

Even when self-describing formats like HDF5 are used, the organization of data—such as the structure, names, and descriptions of storage objects, e.g., groups, datasets or attributes—often still differ between applications and experiments. This diversity makes the development of common and reusable tools for processing, exchange, analysis, and visualization of data challenging. Some formats, e.g., VizSchema [12] and XDMF [3], propose to bridge this gap between general-purpose, self-describing formats and the need for standardized tools for data exchange, processing, and interpretation by augmenting HDF5 via lightweight, low-level schema (often based on XML) to further describe the organization of data. For example, the primary goal of XDMF (eXtensible Data Model and Format) [16, 3] is to help standardize methods to exchange scientific data between high-performance computing codes and tools. XDMF distinguishes and separates so-called light and heavy data. Light data contains the basic description of data arrays—e.g, the value type (float, integer, etc.), precision, location, rank, and dimensions of data arrays— while heavy data refers to the actual multi-dimensional arrays storing scientific data values. XDMF stores light data in XML while heavy data is stored in HDF5. The focus of formats like XDMF and VizSchema is primarily the standardized description of the low-level data organization to facilitate data exchange and tool development.

In contrast to XDMF and VizSchema, application oriented formats generally focus on specifying the organization of data in a semantically meaningful fashion, including but not limited to: the specification of storage object names, locations, descriptions, and data hierarchies. Many scientific application formats build on existing self-describing formats (e.g, HDF5), and examples include the NeXus [8] format for neutron, x-ray, and muon data, the OpenMSI format for mass spectrometry imaging data [11], the CXIDB format [9] for coherent x-ray imaging and many others. Application formats are often described by documents that specify the precise location and names of data items and in many cases provide some form of application-programmer interface (API) to facilitate reading and writing of format files. Some formats are further governed by formal, computer-readable, and verifiable specifications. For example, NeXus uses NXDL2, an XML-based format and schema that allows scientists to define the nomenclature and arrangement of information in a NeXus data file. On the level of HDF5 groups, NeXus also uses the notion of *Classes* to define the fields that a group should contain in a reusable and extensible fashion.

The critical need for data standards in neuroscience research has been recognized by several efforts over the course of the last several years; however, much work remains. Here, our goal is to contribute to this discussion by instantiating a usable and sustainable data standard for neuroscience research. The developers of the *Klustakwik* suite[7, 6] have proposed an HDF5-based data format for storage of spike sorting data. *Orca* (also called *BORG*) is an HDF5-based format developed by the Allen Institute for Brain Science designed to store electrophysiology and optophysiology data ^3^. The *NIX* [13] project has developed a set of standardized methods and models for storing electrophysiology and other neuroscience data together with their metadata in one common file format based on HDF5. Rather than an application-specific format, NIX defines highly generic models for data as well as for metadata that can be linked to terminologies (defined via *od*ML) to provide a domain-specific context for elements. The *open metadata Markup Language od*ML [5] is a metadata markup language based on XML with the goal to define and establish an open and flexible format to transport neuroscience metadata. NeuroML [4] is also an XML-based format with a particular focus on defining and exchanging descriptions of neuronal cell and network models. The neurodata without borders (NWB)^4^ initiative is a recent project with the goal *“[…] to produce a unified data format for cellular-based neurophysiology data based on representative use cases initially from four laboratories – the Buzsaki group at NYU, the Svoboda group at Janelia Farm, the Meister group at Caltech, and the Allen Institute for Brain Science in Seattle.”* Members of the NIX, KWIK, Orca, BRAINformat, and other development teams^5^ have been invited and have contributed to the NWB effort. NWB has adopted concepts and methods from a range of these formats, including from the here-described BRAINformat.

## 3 STANDARDIZING SCIENTIFIC DATA

### 3.1 Data Organization and File Format API

BRAINformat adopts HDF5 as its main storage backend and uses the following primary storage primitives to organize data within files:

- **Group:** A group is used—similar to a folder or directory on a file system—to group zero or more storage objects.
- **Dataset:** A dataset defines a multidimensional array of data elements, together with supporting metadata (e.g., shape and data type of the array).
- **Attribute:** Attributes are small datasets that are attached to primary data objects (i.e., groups or datasets) and are used in practice to store additional metadata to further describe the corresponding data object.
- **Dimension Scale:** This is a derived storage primitive that uses a combination of datasets and attributes to associate datasets with the dimension of another dataset. Dimension scales are used in practice to further characterize the dimensions of a dataset by describing, for example, the time when samples were measured or the location of samples in space.
- **Relationship Attributes:** Relationship attributes are a novel, custom attribute-type storage primitive that allows us to describe and model structural and semantic relationships between primary data objects in a human-readable and computer-interpretable fashion (described later in Section 3.2).

Neuroscience research inherently relies on complex collections of data from many modalities and sources. Examples include neural recordings, audio and video recordings, eye-tracking, motion tracking, task contingencies, external stimuli, derived analytic results, and many others. It is therefore critically important to specify formats in a modular and extensible fashion while enabling users to easily reuse format modules and integrate new ones. The concept of managed objects, which we will describe next, allows us to address this central challenge in an easy-to-use and scalable fashion.

#### 3.1.1 Managed Objects

A managed object is a primary storage object—i.e., file, group, or dataset—with: **1)** a formal, self-contained format specification that describes the storage object and its contents (see Section 3.1.2), **2)** a specific managed type/class, **3)** a human-readable description, and **4)** an optional unique object identifier, e.g., a DOI. In file, these basic managed object descriptors are stored via standardized attributes. Managed object types may be composed—i.e., a file or group may contain other managed objects—and further specialized through the concept of inheritance, enabling the independent specification and reuse of data format components. The concept of managed objects significantly simplifies the file format specification process by allowing larger formats to be specified in an easy-to-manage iterative manner. By encapsulating semantic sub-components, managed objects provide an ideal foundation for interacting with data in a manner that is semantically meaningful to applications.

The BRAINformat library provides dedicated base classes to assist with the specification and development of interfaces for new managed object types. The *M anagedObject* base API implements common features to **1)** define the specification of a given managed type, **2)** recursively construct the complete format specification, automatically resolving nesting of managed objects, **3)** verify format compliance of a given HDF5 object, **4)** provide access to all common managed object descriptors stored in file (i.e., type, description, specification, and object identifier), and provides a standardized interface to **5)** access contained objects (e.g, datasets, groups, managed object etc.) from file, **6)** retrieve all managed object instances of a given managed type, and **7)** create appropriate manager class instances for a given HDF5 object based on the objects managed type.

In addition, the *M anagedObject* base API defines and implements a standardized approach for creation of specific instances of managed objects stored in file via a common *create*(..) method. Managed groups and datasets may be stored either directly within the parent managed group or created externally in a separate *M anagedObjectF ile* file storage container and included in the parent via an external link. In this way, the API directly supports self-contained and modular data storage in a transparent fashion. Self-contained storage eases data sharing, as all data is contained within a single file, while modular storage allows us to more easily manage file sizes and reduce the risk for file corruption by minimizing changes to existing files. From a user’s perspective, modular and self-contained storage are handled transparently, i.e., a user can interact with managed objects in the same manner independent of whether the object is stored internal or external to the current HDF5 file.

To implement a new managed object type, a developer simply needs to define a new class that inherits from the appropriate base managed class type—i.e., *M anagedF ile*, *M anagedGroup*, and *M anagedDataset*—and implement: **1)** the class method *get*_*f ormat*_*specif ication*(*…*) to create a formal format specification document (described next in Sec. 3.1.2) and **2)** the object method *populate*(*…*), which is called by the standardized *M anagedObject.create*(*…*) method and is used to implement the type-specific population of managed storage objects to ensure format compliance upon creation—i.e., the goal is to avoid that managed objects can be created in an invalid, non-format-compliant state to ensure that files remain format compliant throughout their life cycle.

#### 3.1.2 Format Specification

To enable the broad application and use of data formats, it is critical that the underlying data standard is easy to interpret by application scientists as well as unambiguously specified for programmatic interpretation and implementation by developers. Therefore, each data format component (i.e, managed object type) is described by a formal, self-contained format specification that is computer interpretable while at the same time including human-readable descriptions of all components.

We generally assume that format specifications are minimal, i.e., all file objects that are defined in the specification must adhere to the specification, but a user may add user-defined data objects (i.e., groups, datasets, attributes etc.) to a file without violating format compliance. The relaxed assumption of a minimal specification ensures on the one hand that we can share and interact with all format-compliant files and file components in a standardized fashion, while at the same time enabling users to easily integrate dynamic and custom data (e.g, instrument-specific metadata), allowing researchers to save all their data using BRAINformat even if the current file standard should only partially cover the specific use-case. This is critical to enable scientists to easily adopt the file standard and to allow the file standard to adapt to the ever-evolving experiments, methods, and use-case in neuroscience and facilitate new science rather than impeding it.

The BRAINformat library defines format specification document standards for the specification of the format of **1)** files, **2)** groups, **3)** datasets, **4)** attributes, **5)** dimension scales, **6)** managed objects, and **7)** relationship attributes. All specification documents are based on hierarchically composed Python dictionaries that can be serialized as JSON documents for persistent storage and sharing. For all data objects we specify the name and/or prefix of the object, whether the object is optional or required, and provide a human-readable textual description of the purpose and content of the object. Depending on the object type (e.g, file, group, dataset, attribute, etc.) additional information is specified, e.g., i) the datasets, groups, and managed objects contained in a group or file, ii) attributes for datasets, groups and files, iii) dimension scales of a dataset, iv) whether a dataset is a primary dataset for visualization and analysis or iv) relationships between objects among others. Figure 1 shows as an example an abbreviated summary of the format specification of our proposed data standard for neuroscience (described later in Section 4). Supplement 2 provides a more detailed discussion and examples of our format specification model. Relationship attributes and their specification are discussed in detail later in Sec. 3.2.

**Figure 1.**
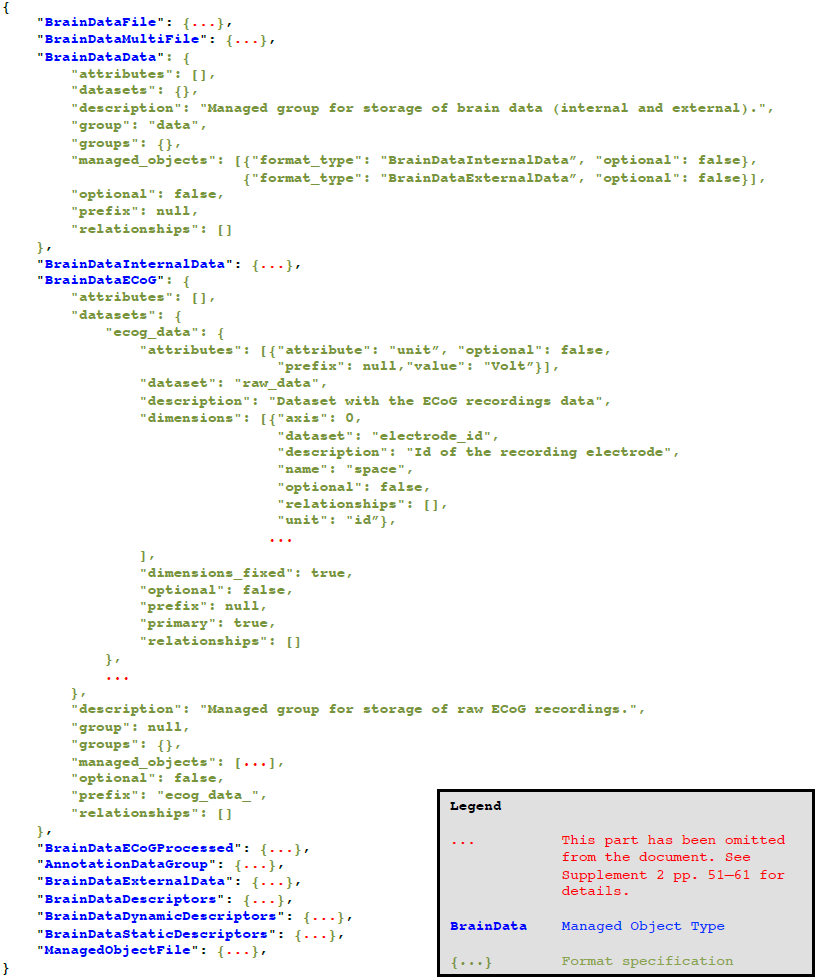
Abbreviated specification document for our neuroscience data format listing all current managed object types and partial specification for select structures illustrating the general structure of a formal specification document generated using the BRAINformat library. The full specification document is available as part of Supplement 2 pp. 51 – 61 (and the full recursive specification for a brain format file is shown in Supplement 2 pp. 35 – 51).

The BRAINformat library implements a series of dedicated data structures to assist with the development and interaction with format specifications. Using the provided data structures helps ensure that the generated documents are valid—e.g., that all required keys are set and that only valid keys and values are included in a document—and supports the incremental creation of format specifications, allowing the developer to step-by-step define and compose format specifications—similar to how one typically creates HDF5 files. For example, the following simple code can be used to generate the parts of the *BrainDataECoG* specification shown in Fig. 1:

~~~
>>> from brain.dataformat.spec import *
~~~

~~~
>>> **# Define the raw dataset and associated attribute and dimension**
~~~

~~~
>>> raw_data_spec = **DatasetSpec(**dataset=’raw_data’, prefix=None, optional=False, primary=’True’,
~~~

~~~
description="Dataset with the ECoG recordings data")
~~~

~~~
>>> raw_data_spec.add_attribute (**AttributeSpec(**attribute=’unit’, prefix=None, value=’Volt’))
~~~

~~~
>>> raw_data_spec.add_dimension (**DimensionSpec(**name=’space’, unit=’id’, dataset=’electrode_id’,
~~~

~~~
axis=0, description="Id of the recording electrode"))
~~~

~~~
>>> **# Define the group and add the dataset**
~~~

~~~
>>> brain_data_ecog = **GroupSpec(**group=None, prefix=’ecog_data_’,
~~~

~~~
description="Managed group for storage of raw ECoG recordings.")
~~~

~~~
>>> brain_data_ecog.add_dataset(raw_data_spec, ’ecog_data’)
~~~

Using the BRAINformat specification infrastructure we can easily compile a complete data format specification document that lists all managed object types and their format. For example, the simple Python code shown here compiles the format specification document for our neuroscience data format directly from the Python API of our format (see also Sec. 4):

~~~
>>> from brain.dataformat.spec import FormatDocument
~~~

~~~
>>> import brain.dataformat.brainformat as brainformat
~~~

~~~
>>> **json_spec = FormatDocument.from_api(module_object=brainformat).to_json()**
~~~

Figure 1 shows an abbreviated summary of the result of the above code. The full JSON document is shown in Supplement 2, pp. 51 – 61. Alternatively, we can also recursively construct the complete specification for a given managed object type —e.g., here for the main file of the proposed neuroscience format described in Sec. 4— via:

~~~
>>> from brain.dataformat.brainformat import BrainDataFile
~~~

~~~
>>> from brain.dataformat.spec import *
~~~

~~~
>>> **format_spec = BrainDataFile.get_format_specification_recursive()** # Construct the document
~~~

~~~
>>> **file_spec = BaseSpec.from_dict(format_spec)** # Verification of the document
~~~

~~~
>>> **json_spec = file_spec.to_json(pretty=True)** # Convert the document to JSON
~~~

In this case, all references to other managed objects are automatically resolved and their specification is directly embedded in the resulting specification document. While the basic specification for *BrainDataF ile* consists only of *≈*14 lines of code (see Supplement 2, pp. 30), the full, recursive specification contains more than 910 lines (see Supplement 2, pp. 35 – 51), illustrating the critical importance for being able to incrementally define format specifications.

The ability to compile complete format specification documents directly from data format APIs allows developers to easily integrate new format components (i.e. managed object types) in a self-contained fashion simply by adding a new API class without having to maintain separate format specification documents. Furthermore, this strategy avoids inconsistencies between data format APIs and specification documents since format documents are updated automatically.

The concept of managed objects in combination with the format specification language and API provide an application-independent design concept that allows us to define application-specific formats and modules that are build on best practices.

### 3.2 Modeling Data Relationships

Neuroscience data analytics often rely on complex structural and semantic relationships between datasets. For example a scientist may use audio recordings to identify particular speech events during the course of an experiment and in turn needs to locate the corresponding data in an electrocorticography recording dataset to study the neural response to the speech events. In addition, we often encounter structural relationships in data, for example, in the case of data structures where one array indexes another array or two arrays share data dimensions because they have been acquired using the same recording device and many others. To enable efficient analysis, reuse, and sharing of neuroscience data it is critical that we can model the complex relationships between data objects in a structured fashion to enable human and computer interpretation and use of data relationships.

Modeling data relationships is not well-supported by traditional data formats, but is typically closer to the domain of scientific databases. In HDF5, we can compose data via HDF5 links (soft and hard) and associate datasets with the dimensions of another dataset via the concept of dimension scales. However, these concepts are limited to very specific types of data links that do not describe the semantics of the relationship. A new general approach is needed to describe more complex semantic links between data objects in HDF5.

#### 3.2.1 Specifying and Storing Relationships

Here we introduce the novel concept of *relationship attributes* to describe complex semantic relationships between a source object and a target data object in a general and extensible fashion. Relationship attributes are associated with the source object and describe how the source is related to the target data object. The source and target of a relationship may be either a HDF5 group or dataset.

Relationship attributes are—like other file components— specified via a JSON dictionary and are part of the specification of datasets and groups. Like any other data object, relationships may also be created dynamically to describe any relationships that are unknown *a priori*. Specific instances of relationships are stored as attributes on the source HDF5 object, where the value of the attribute is the JSON document describing the relationship. As illustrated in Fig. 2, the JSON specification of a relationship consists of the following main components:

**Figure 2.**
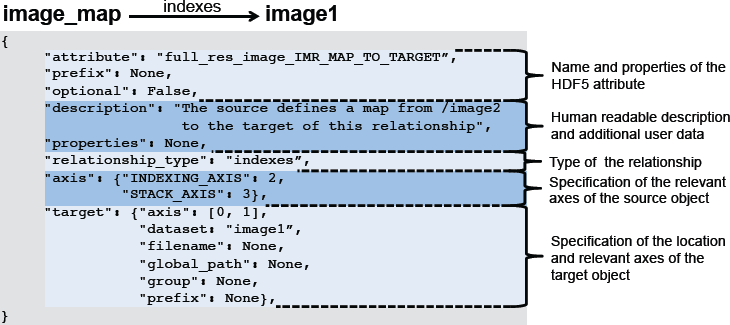
Example specification of a relationship attribute illustrating the main components of the specification.

1. The specification of the name of the attribute and whether the attribute is optional. When stored in HDF5 we prepend the prefix *RELATIONSHIP* _*AT T R*_ to the user-defined name of the attribute to describe the attribute’s class and ease identification of relationship attributes.
2. A human-readable description of the relationship and an optional JSON dictionary with additional user-defined data relevant to the relationship.
3. The specification of the type of the relationship (described next in Sec. 3.2.2).
4. The specification of the axes of the source object to which the relationship applies. This may be: i) a single index, ii) a list of axes, iii) a dictionary of axis indices if the axes have a specific user meaning, or iv) None if the relationship applies to the source object as a whole. Note, we do not need to specify the location of the source object, as the specification of the relationship is always associated with either the source object in HDF5 itself or in the format specification.
5. The specification of the target object describing the location of the object and the axes relevant to the relationship (using the same relative ordering or names of axes as for the source object).

#### 3.2.2 Relationship Types

The relationship type describes the semantic nature of the relationship. The BRAINformat library currently supports the following main types of relationships, and additional types can be added in the future:

- **order:** This relationship type indicates that elements along the specified axes of the relationship are ordered in the target in the same way as in the source. This type of relationship is very common in practice. For example, in the case of dimension scales, an implicit assumption is that the ordering of elements along the first axis of the scale-dataset matches the ordering of the elements of the dimension it describes. This assumption, however, is only implicit and is by no means always true (nor does HDF5 require this relationship to be true). Using an *order* relationship we can make this relationship explicit. Other common uses of *order* relationships include describing the matched ordering of electrodes in datasets that have been recorded using the same device or matched ordering of records in datasets that have been acquired synchronously.
- **equivalent:** This relationship type expresses that the source and target object encode the same data (even if they might store different values). This relationship also implies that the source and target contain the same number of values ordered in the same fashion. This relationship occurs in practice any time the same data is stored multiple times with different encodings. For example to facilitate data processing a user may store a dataset of strings with the names of tokens and store another dataset with the equivalent integer ID of the tokens.
- **indexes:** An indexes relationship describes that the source dataset contains indices into the target data object (group or dataset). In practice this relationship type is used to describe basic data structure where we store, for example, a list of unique values (tokens) along with other arrays that reference that list.
- **shared encoding:** This relationship indicates that the source and target data object contain values with the same encoding so that the values can be directly compared (via equals "=="). This relationship is useful in practice any time two data objects (datasets or groups) contain data with the same encoding (e.g. two datasets describing external stimuli using the same ontology).
- **shared ascending encoding:** This relationship type implies that the source and target data object share the same encoding and in addition that the values are sorted in ascending order in both data objects. The additional constraint on the ordering enables i) comparison of values via greater than "*>*" and less than "*<*" (in addition to equals ==) and ii) more efficient processing and comparison of data ranges. For example, in the case of two datasets that encode *time*, we often find that individual time points do not match exactly between the source and target (e.g, due to different sampling rates). However, due to the ascending ordering of values, a user is still able to compare ranges in *time* in a meaningful way.
- **indexes values:** This relationship is typically used to describe value-based referencing of data and indicates that the source data object selects certain parts of the target data object based on data values (or keys in the case of groups). This relationship is a special type of *shared encoding* relationship.
- **user:** The *user* relationship is a general container to allow users to specify custom semantic relationships that do not match any of the existing relationship patterns. To further characterize the relationship, we often store additional metadata about the relationship as part of the user-defined *properties* dictionary of the relationship attribute.

#### 3.2.3 Using Relationship Attributes

Relationship attributes are a direct extension to the previously described format specification infrastructure. Similar to other main data objects, BRAINformat provides dict-like data structures to help with the formal specification of relationship attributes. In addition, the BRAINformat library also provides a dedicated *RelationshipAttribute* API, which supports creation and retrieval of relationship attributes (as well as index map relationship, described in Sec. 3.2.4) and provides easy access to the source and target HDF5 object and corresponding specifications of relationship attributes.

One central advantage of explicitly defining relationships is that it allows formalizing the interactions and collaborative usage of related datasets. In particular, the relationship types imply formal rules for how to map data selections from the source object of a relationship to the target object. The *RelationshipAttribute* API implements these rules and supports slicing, which allows us to easily map selections from the source to the target data object using the same familiar slicing syntax. For example, assume we have two datasets *A* and *B* related to each other via an *indexes* relationship *R*_*A→B*_. A user now selects the values *A*[1 : 10] in the source dataset *A* and wants to locate the corresponding data values in the target data object *B*. Using the BRAINformat API we can now simply write *R*_*A→B*_[1 : 10] to map the selection [1 : 10] from the source *A* to the target *B*, and if desired retrieve the corresponding data values in *B* via *B*[*R*_*A→B*_[1 : 10]]. Figure 3 provides an overview of the rules for mapping selections based on the type of the relationship.

**Figure 3.**
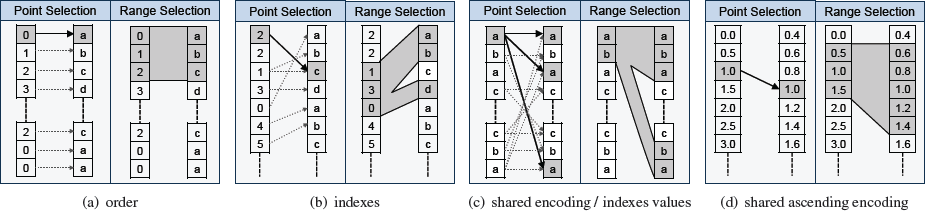
Overview of the main relationship types and the implied mapping of point-and range-based selections from the source to the target object. In each cell we show the source object on the left and the target object of the relationship on the right. **(a)** For *order* relationships we can directly map array indices between the data objects. In the case of *order* relationships involving HDF5 Groups we assume alphabetic ordering of elements. **(b)** In the case of *indexes* relationships we map selections by retrieving the relevant indicies from the source array. **(c)** For *shared encoding* and *indexes values* relationships we support data selection via value-based data mapping, i.e., we map selections by locating all data values in the target object that match at least one of the values we selected in the source object. **(d)** *Shared ascending encoding* relationships behave in general similar to *shared encoding* relationships, however, the additional constraint that values are sorted in ascending order enables us to map range selections directly based on the minimum and maximum value selected in the source dataset (in contrast to the strict equal value matching of *shared encoding*). **(e)** *User* relationships define custom user semantics and do not imply a specific mapping between data elements (not shown).

Relationship attributes standardize the specification, storage, and programmatic interface for creating, discovering, and using relationships and related data objects. Describing relationships between data explicitly greatly simplifies the process of interacting with multiple datasets and facilitates the collaborative use of data by enabling utilization of multiple datasets in conjunction without having to *a priori* know the relationships and datasets involved. In this way, relationship attributes also open the route for the standardized development of novel data-driven analytics and workflows based on the programmatic discovery and use of related data objects.

#### 3.2.4 Index Map Relationships

Beyond the description of direct object-to-object relationships, relationship attributes also form the building blocks that allow us to specify higher-order relationships. Using relationship attributes we can define chains of object-to-object relationships that, when interpreted in conjunction, express highly complex structural and semantic relationships. For example, imagine the following situation. Scientists have acquired an optical microscopy image *A* and an electron microscopy image *B* of the same brain. Using the optical image a scientist identifies a particular brain region of interest and now wants to study the same region further using the electron-microscopy image. This seemingly simple task of accessing corresponding data values in two related datasets is in practice, however, often highly complex. Even if the data registration problem between the datasets is solved, a user still has to know exactly: i) the location of both datasets *A* and *B*, ii) how the two datasets are related, iii) what the transformations generated by the data registration process are, iv) how to utilize that information to map between *A* and *B*, and v) write complex, custom code to access the data.

*Index map relationships* allow us to explicitly describe this complex relationship between *A* and *B* via a simple chain of object-to-object relationship attributes and to greatly simplify the cooperative interaction with the data. Rather than describing the relationship between *A* and *B* directly, users can create an intermediate index map *M*_*A→B*_ that stores for each pixel in *A* the index of the corresponding pixel(s) in *B*. *M*_*A→B*_ explicitly and unambiguously describes the mapping from *A* to *B* so that we can directly utilize the mapping without having to perform complex and error-prone index transformations (which would be needed if we described the mapping implicitly, e.g., via scaling, rotation, morphing and other data transformations). As the table in Fig. 4 shows, via a simple series of relationship attributes describing simple object-to-object relationships, we can unambiguously describe the complex relationship between *A* and *B* via *M*_*A→B*_. Given only our source dataset *A* (or index map *M*_*A→B*_) we can now easily discover all relevant data objects (*A*, *B*, and *M*_*A→B*_) and relationships (Fig. 4) without having to *a priori* know the mapping or the location of the datasets. Via the index map relationship, we can now directly map selections: i) from *A* to *M*_*A→B*_ and *vice versa* ii) from *M*_*A→B*_ to *B*, and most importantly iii) from *A* to *B* simply by slicing into our *indexes* relationship (Fig. 4, row 3) to retrieve the corresponding indices from our index map *M*_*A→B*_. As data mappings are described explicitly, index map relationships enable registration and mapping under arbitrary transformations. Also, mappings are not required to be unique—i.e., arbitrary N-to-M mappings between elements are permitted—and the source and target of relationships may not just be datasets but also groups, i.e., index map relationship can be used to define mappings between contents of groups or even groups and datasets in HDF5.

**Figure 4.**
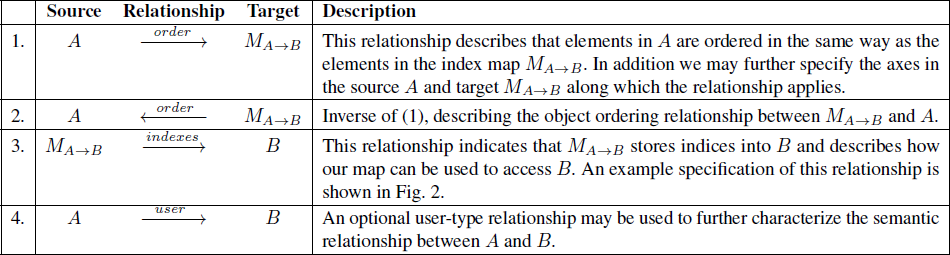
Overview of the relationships used to define an advanced *index map relationship*. We present a specific example later in Fig. 5.

**Figure 5.**
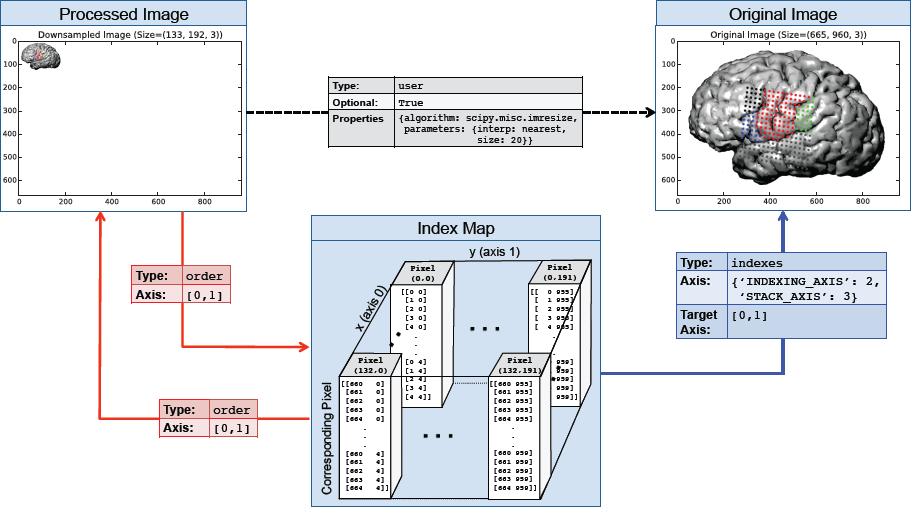
Illustration of an index map relationship describing the interaction between a processed image and the original image. The processed image is in this case a 5*×* smaller version of the original image created using nearest neighbor interpolation. The intermediary index map describes for each pixel in the processed image which pixels it corresponds to in the the original image. Two *order* relationships (red arrows) describe the interactions between the processed image and the map and vice versa. A third *indexes* relationship links our index map to the original image and describes how the map can be used to access the original image. Optionally, we may create a fourth *user* relationship (black arrow) to further characterize the semantic relationship between the processed and original image (e.g, to store a description of the algorithm and parameters used to generate the image). Naturally, we can also describe the inverse mapping between the original and processed image via a second index map relationship.

BRAINformat implements the concept of index map relationships—similar to dimension scales and relationship attributes— via a set of simple naming conventions for the attribute names. In addition to the *RELATIONSHIP_ATTR* prefix, we use a set of reserved post-fix values—specifically *_IMR_MAP_TO_TARGET*,*_IMR_MAP_TO_SOURCE*, *_IMR_SOURCE_TO_MAP*, *_IMR_SOURCE_TO_TARGET*—that are appended to the user-defined attribute name to identify the different components of the index map relationship. The BRAINformat API directly supports index map relationships so that we can, for example, directly create and locate all relationships that define an index map relationship via a single function call and programmatically interact with the relationships. Supplement 1 (pp.12–26) includes an overview and basic tutorial of the API for creating and using index map relationships.

Index map relationships have broad practical applications, including data registration, sup-component analysis, correlation of data dimensions, and optimization. Index map relationships are directly applicable to specify the mapping between images in a time series or a stack of physical slices as well as to define correspondences between images from different modalities. We may also define mappings between select dimensions of a dataset to correlate data from different recordings in time or space. Furthermore, analytics are often based on characteristic sub-components of a dataset. As such, a user may extract and separately process sub-components of datasets (e.g. a sub-image of a single cell) and use index map relationships to map the extracted or derived analysis data back to the original data. To optimize data classification, feature detection, and other compute-intensive analyses, a user may perform initial calculations on lower-resolution versions of a dataset and use index map relationships to access corresponding data values in the high-resolution version of the dataset for further processing.

Figure 5 illustrates an example index map relationship for the latter use-case. The complete source code and further details for this example are available in Supplement 3. In this example, our original dataset is a RGB image dataset of size (665 *×* 960) that has been processed to reduce the size in the two spatial dimensions by a factor of 5 to (133 *×* 3) via nearest neighbor interpolation. Each pixel in the processed image, hence, maps to a 5 *×* 5 sub-region in the original image. We, hence, create a 4-dimensional index map dataset where: 1,2) the first two dimensions correspond to the spatial dimensions *x* and *y* of the images, 3) the third dimension is our index axis of length 2 since each pixel is described by two integer indices, and 4) the fourth dimension is our stacking axis with the list of all corresponding pixel. Using the BRAINformat API, we can now create the index map relationship—which is defined by the arrows shown in Figure 5—via a single function call (Supplement 3–Sec. 1.3). As illustrated in Figure 6, we can now easily map a selection (here [47, 98]) from our source (processed image) to the target (original) image simply by slicing into our index map relationship (*imr*) via *imr*[*M AP* _*T O*_*T ARGET* ][47, 98] and retrieve the data of the corresponding subimage from our original image (Supplement 3–Sec. 1.4).

**Figure 6.**
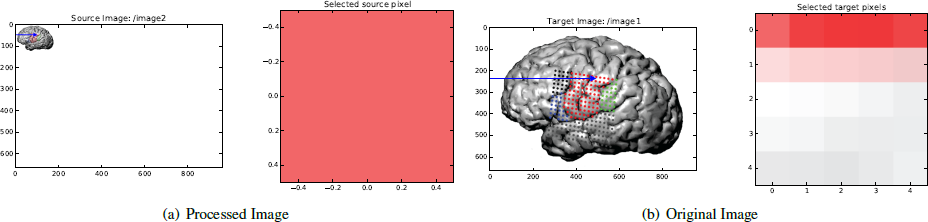
Illustration of the result from using our index map relationship to perform data selection in our source, processed image (a) and target, original image (b). **(a)** First we apply the selection (47, 98) (blue arrow) to our source dataset (left). As expected, this results in the selection of a single pixel (right). **(b)** We next map the same selection to our target dataset (left). From the blue arrow we can see that the selection was mapped correctly to same relative location as in our source, image. The pixel plot (right) illustrates that the mapping resulted, as expected, in the selection of a 5 *×* 5 sub-image from our target image. We can also see that the top-left pixel of our selected sub-image matches the color of the pixel we retrieved in the source dataset (a, right). This is expected since the source image (a, left) was generated from the target image (b, left) via 5*×* downsampling using nearest neighbor interpolation.

As this simple example illustrates, index map relationships allow us to explicitly describe complex relationships between data. Being able to unambiguously describe complex relationships is critical to enable us to programmaticaly utilize relationships and perform complex multi-data analytics and to reduce risk for errors due to implicit assumptions about relationships between data objects. Index map relationships are not restricted to just define relationships between HDF5 datasets but can also be used to define relationships involving HDF5 groups or managed objects. Here we focus on index map relationships, but the same basic concept of chaining relationships could be applied to construct other types of complex object inter-relationships as well.

### 3.3 Annotating Scientific Data

Advanced neuroscience analytics rely on complex data access patterns driven by data semantics. For example, common neuroscience data analytics often focus on understanding how different brain regions—measured, e.g., by collocated electrodes— operate together and interact with each other during specific, randomly interleaved events, e.g., time intervals when a subject said *‘baa‘* or performed a particular motion. The ability to annotate data by associating semantic metadata with data subsets is critical to facilitate these kinds of analyses. Annotating data in a scalable and usable fashion is challenging and relies on complex data structures to describe data selections and associated metadata.

To support data annotation, the BRAINformat library provides a series of modules that implement general and reusable data structures to describe individual data selections and data annotations (Sec. 3.3.1) and modules to manage and store collections of data annotations (Sec. 3.3.2). The BRAINformat annotation package supports annotation of in-memory data arrays (e.g., numpy arrays) as well as in-file arrays (i.e., HDF5 datasets) and processing of data annotations may be performed in-memory or out-of-core (i.e., with the majority of data residing on disk and being only loaded when needed).

#### 3.3.1 Data Selection and Annotation

The first steps in annotating data is to describe 1) the data object that contains the data and 2) the data selection describing the subset of the data to annotate. The first part of describing the data object itself is generally simple and consists of either a basic reference to the data object in memory or an HDF5 link to the corresponding object in file. Describing data selections, however, is in practice not as simple. In the context of neuroscience data, researchers often need to generate a large numbers of annotations that refer to complex subsets of data, leading to advanced data selection, storage, and API requirements for describing, storing, and interacting with data selections. For example, along a single axis (such as time), features of interest are often discontinues—e.g., when describing multiple events of the same type—and complex features are often the result of combinations of basic features along multiple dimensions—e.g., the output measured by electrodes located in the hippocampus while the animal is in a specific location.

The table in Fig. 7 provides a high-level comparison of four common schema for describing data selections (columns) with respect to their general behavior in regard to some main requirements for annotating neuroscience data (rows). Slicing is a very convenient way to express highly structured selections that can be described via a simple tuple of (*start, stop, step*) but it does not support selection of complex data subsets. Binary vectors—describing for each element along a given axis whether the element is selected—are generally a good option. One main disadvantage of binary vectors is that the memory cost can be high when having to process a large number of selections in uncompressed form in memory. In practice, however, most operations can be performed iteratively and out-of-core. Lists of indices—describing along each axis the specific, selected elements—are also a very good option. The main disadvantage of index lists lies in the high cost for describing dense selections and the variable length arrays needed to describe the selections. More advanced data selection methods, such as, word-aligned hybrid compressed bitmap indices [17, 18] are also very promising. One main disadvantage of such advanced indexing schema is that they are not easily interpreted without a dedicated API, potentially hindering reuse of the HDF5 files. For the initial development of the BRAINformat annotation API and format we have chosen binary vectors as the main scheme to represent complex data selections and are planing to add support for additional schema in the future.

**Figure 7.**
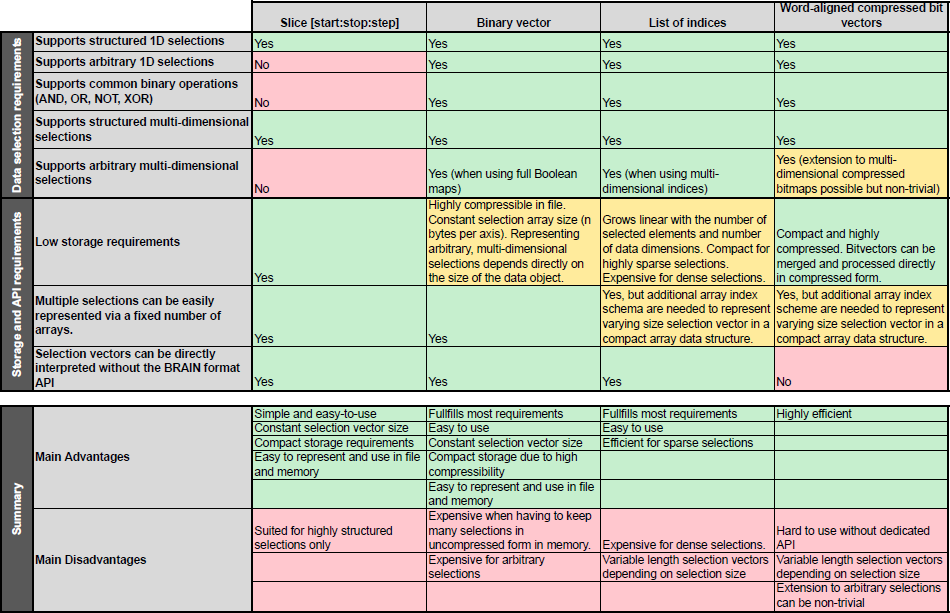
High-level comparison of four common schema for representing data selections. In each case, structured, multi-dimensional selections are constructed by the intersection (AND) of one-dimensional selections along the individual axes while *None* is used to efficiently describe the selection of all elements along a given axis.

Fig. 8(a) illustrates how we can represent and combine complex selections using binary vectors. Along each dimension we store a binary vector describing the elements that are selected (True, color) or not selected (False, white). This allows us to easily represent arbitrary selections using constant-length selection vectors. The binary vectors can be efficiently combined directly using common binary operations and multi-dimensional selections can be easily described by the intersection of multiple binary vectors. Arbitrary multi-dimensional selections—needed to describe complex, multi-dimensional combinations of basic selections and arbitrary user-defined selections—can be described via full binary selection masks (see Fig. 8(b)).

**Figure 8.**
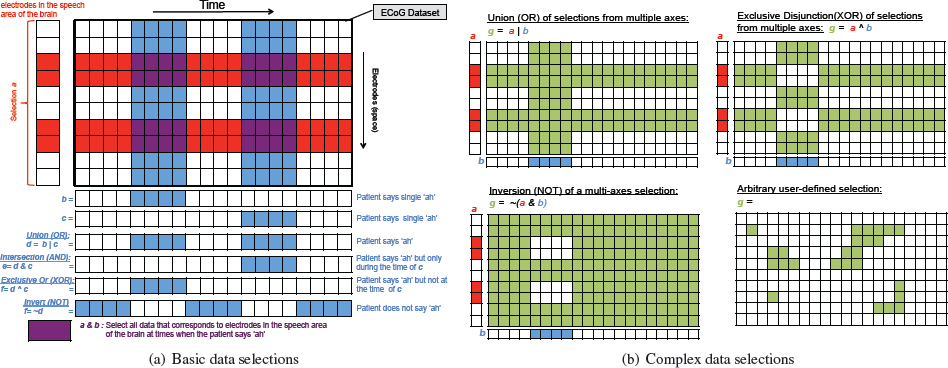
Overview of common data selections and binary combinations of data selections. **(a)** Basic data selections are defined via 1D binary vectors demarking the elements selected along a given dimension. Basic multi-dimensional selections are then defined via binary *AN D* (&) combinations of such per-axis binary vectors. This allows complex selections to be expressed and stored efficiently. Along a given axis we may combine selections via boolean operations—including, *AN D*, *OR*, *XOR*, and *N OT* —without the need to expand the selection to a complex selection. **(b)** We support complex selections that cannot be expressed via combinations of hyper-slices through expansion of the selection to a full binary map allowing the definition of arbitrary selections. Complex selections are required in practice to define *OR*, *XOR*, and *N OT* combinations of multi-dimensional selections and for arbitrary user selections.

Now that we can describe data selections, we can extend our design to define annotations. A single data annotation in BRAINformat consists of the following main elements: **i)** the data selection object describing the data object and subregion of the data the annotation applies to, **ii)** a user-defined string indicating the type of the annotation, **iii)** a human-readable description of the annotation, and **iv)** a dictionary of additional user-defined properties of the annotation, Currently the format requires that the keys of the properties dictionary are strings and that the values are arbitrary, basic data objects, e.g, stings or numbers. This simple design allows us to describe complex annotations in an easy-to-use fashion.

The BRAINformat data selection and annotation API can be used to annotate any data object that can describe its shape as an n-dimensional array and supports numpy/h5py-style array slicing, including numpy arrays, HDF5 datasets, and certain managed objects that implement an array-like interface. The data selection and annotation API supports (among other things):

- Selection of elements via basic array slicing and assignment, e.g., to select the first five elements along the *time* axis for a data selection *A*, we may write *A*[*′time′,* 0 : 5] = *True*.
- Retrieval of the selection vector along a given axis via simple slicing, e.g, *A*[*′time′*].
- Retrieval of the data selected by an annotation or data selection via *A.data*().
- Common binary operations to merge data selections and annotations, including i) AND, ii) OR, iii) NOT, and iv) XOR.
- Comparison of data selections and annotations via common operations, such as *>*, *>*=, *<*, *<*=, ==, ! =, and *in*. These operations are based on the comparison of the selected array indices so that, e.g., *A > B* is only true if *B* is a true subset of
- *A* (in contrast to a simple length-based comparison which would only require *|A| > |B|*).
- Preceded (*A << B*) and follows (*A >> B*) operations describing whether all elements selected by *A* have array indices less than or greater than *B*, respectively. This is useful, for example, to identify if an event in *time* selected by *A* occurs before/after *B*.
- Basic investigation via functions like *len*, *count*, *counts*, *axes* or *axis*_*bounds* to retrieve the total and per-axis number of elements selected or the axes that are restricted and index ranges *et cetera*.

#### 3.3.2 Managing Collections of Data Annotations

So far we have focused on describing single annotations. Neuroscience analytics often rely on large collections of annotations to describe, e.g., multiple behavioral measures, external stimuli, brain regions and many other types of annotations. During the course of an experiment we often encounter many thousands of events and features of a given type and in some cases millions (e.g, action potentials). Storing all these annotations individually would quickly result in an explosion of datasets and groups in the HDF5 file, hindering both usability and computational performance. It is, therefore, critical that we can represent collections of annotations in a compact fashion via a limited number of data objects (Fig. 7). In addition, we need to be able to easily search collections of annotations to locate subsets of annotations of interest, e.g., all annotations describing movements to a specific region in space.

Annotation collections are managed in the BRAINformat library by the *AnnotationDataGroup* (and *AnnotationCollection*) module, which uses the *M anagedObject* design (see Section 3.1.1) to define a general, reusable, and extensible storage module and format for collections of data annotations. Each collection of annotations applies to a specific data object and has a user-defined description. The corresponding binary selection vectors of all annotations in a given collection have the same length, since all annotations refer to the same data object. For each data axis, we can store an arbitrary number of selections in a single two-dimensional data array with a shape of #*selections ×* #*values*. To reduce storage cost, we enable *gzip* compression—which is natively supported by HDF5—when saving the binary data selection arrays to file. Using compression drastically reduces the size of data selections in file and enables us to efficiently store large collections of data annotations. The type, descriptions, and individual properties of all annotations are then stored separately in one-dimensional arrays. This simple scheme allows us to store an arbitrary number of annotations in a fixed number of arrays while allowing us to easily retrieve specific annotations as well as independently access individual fields for searching (e.g., annotation type, description, and individual properties).

Collections of annotations may be created in-memory or in-file. When accessing collections of annotations that are stored in-file, the bulk of the data—such as the selection properties and binary selection vectors—typically remain out-of-core and are only read when needed,for example when searching for annotations based on a particular property. To ease the use of collections of annotations, the BRAINformat API supports:

- **Filtering** (i.e., search) of annotations to locate annotations based on the:
  i. index of annotations,
  ii. axes that are restricted by the annotations to find, e.g., all annotations that select features in *time*,
  iii. full or partial type of annotations to find, e.g., all annotations that define a *speech event*,
  iv. full or partial description of annotations, and
  v. full or partial user-defined metadata properties of annotations to find e.g., all annotations that have a particular user-level, start time or stop time etc..
  All filter functions return one-dimensional binary selection vectors that can be easily combined via standard binary operations. This allows us to easily define complex queries. For example to locate all *speech events* when a subject said *’baa’* in a collection of annotations *C*, we can simply write *C.type*_*f ilter*(*′speech event′*) & *C.property*_*f ilter*(*key* =*′vocalization′, value* =*′ baa′*).
- **Selection** of subsets of annotations, i.e., the creation of new collections of annotations through the application of a filter via basic slicing, e.g., *C*[ *C.type*_*f ilter*(*′speech event′*)].
- **Merging** of all annotations in a collection to a single annotation via union (*OR*), intersection (*AN D*), and exclusive disjunction (*XOR*).
- **Loading/retrieval** of the individual annotations and data selected by the annotations in the collection.
- **Basic introspection** to retrieve information about, e.g., the number of annotations, list of unique annotation types and descriptions, properties, etc..
- **Creation, expansion, and saving** of annotation collections.

Other more specialized functions of annotation collections include the calculation of a containment matrix, describing which annotations are contained in each other. This is useful in the context of hierarchically organized annotations. For example, in the context of speech, we have annotations that describe individual *syllables*, *words*, *sentences* and so on.

## 4 RESULTS

### 4.1 Applications to Neuroscience Data

In the following we describe the application of the BRAINformat data standardization framework to the development of a data format for neuroscience data with an initial focus on electrocorticography (ECoG) data collected from neurosurgical patients during speech production. This data shares many requirements with standard electrophysiology data collected by the broader neuroscience community: storage of voltage recordings over time across multiple spatial distributed sensors with heterogenous geometries, complex and multi-tiered task descriptions, post-hoc processing of raw data to extract the signal of interest, the association of physiology data with multi-modal data streams collected simultaneously by other devices, the linkage of data associated with the same ‘task’ across multiple sessions, and the necessity to store rich meta-data to make sense of it all.

#### 4.1.1 High-level Data Organization

Fig. 9 shows and example visualization of a BRAINformat file using HDFView. The tree view shown on the left illustrates the high-level data hierarchy. In our discussion of the high-level data organization we use the following notation to denote the path in HDF5 and corresponding managed type: ***path : managed type***.

**Figure 9.**
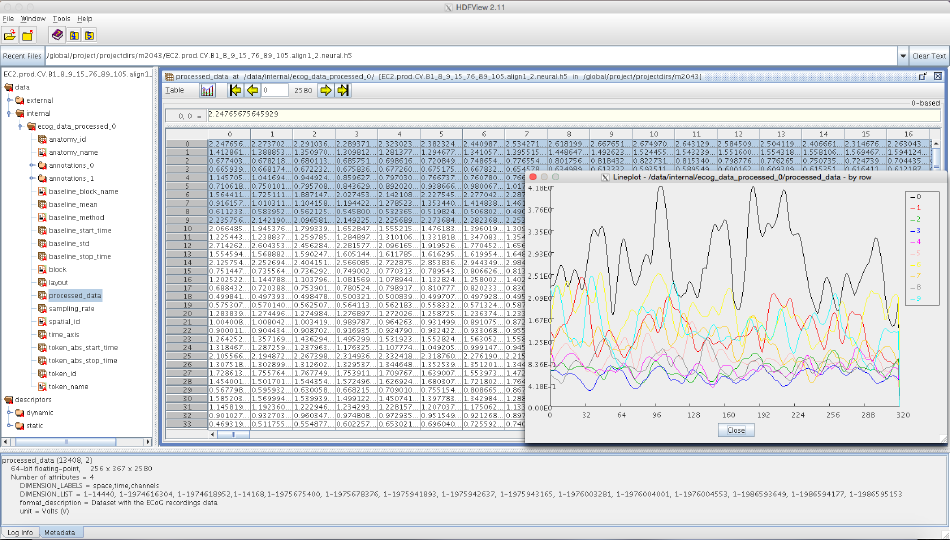
Example HDFView [15] visualization of a BRAINformat file. **Left:** Tree view of the basic data hierarchy. **Right:** Table view of a processed ECoG dataset and curve plot of the first 10 waveforms. **Bottom:** Summary of properties and attributes of the processed ECoG data array.

In the main HDF5 file ***/ : BrainDataFile*** the data is organized in a basic semantic hierarchy. On the highest level we distinguish between data and descriptors, i.e., raw and processed data generated through experimentation and analysis vs. globally accessible metadata. We then further distinguish between static metadata (i.e. descriptions of the basic data acquisition and experimental parameters) and dynamic metadata (e.g. descriptions of post-processing parameters) and categorize data into internal data (i.e. data collected inside the brain, e.g. electrophysiology recordings) and external data (i.e. data collected external to the animal, e.g. sensory stimuli, audio recordings, position of body parts, etc.). These divisions are not strictly necessary, but impose some minimal structure on the format that eases the interpretability by users. The following list illustrates the high-level data organization in more detail:

- ***/data : BrainDataData*** contains the actual raw and processed data generated through experimentation and analysis.
- ***/data/internal : BrainDataInternalData*** contains all internal data, i.e., all raw and processed data from physiological measurements.
- ***/data/internal/ecog_data_# : BrainDataECoG*** is designed for storage of voltage recordings over time across multiple spatially distributed sensors with heterogenous geometries, and complex and multi-tiered task descriptions (see Sec. 4.1.2).
- ***/data/internal/ecog_data_processed_#:BrainDataECoGProcessed*** is derived from ***BrainDataECoG*** and is designed for storage of post-processed voltage recordings over time across multiple spatial distributed sensors where signals of interest have been extracted and optionally categorized (see Sec. 4.1.3).
- ***/data/external : BrainDataExternalData*** is used to collect all external data, such as recordings of sensory stimuli and other external measurements.
- ***/descriptors : BrainDataDescriptors*** is a container for global metadata. Specific metadata objects may be referenced in other managed objects via HDF5 links. This strategy avoids redundant storage while at the same time providing easy access to the data from specific data groups and allowing scientists to collect general metadata in a central location, facilitating meta-and mega analysis.
- ***/descriptors/static : BrainDataStaticDescriptors*** is a container for static metadata, e.g., metadata describing the instruments and other fixed information.
- ***/descriptors/dynamic : BrainDataDynamicDescriptors*** is a container for dynamic metadata, e.g., information that is derived through post-hoc analyses or metadata that may dynamically change during the data life cycle.

In practice, scientists regularly acquire data in series of distinct experiment sessions often distributed over long periods of time. To facilitate management and sharing of data, it is useful to store the data generated from such distinct recordings in separate data files, yet for analysis purposes the data often needs to be analyzed in context. To allow the organization of related data files we support the grouping of files in container files ***/ :BrainDataMultiFile*** in which each primary ***BrainDataFile*** file is represented by an HDF5 group ***/entry_#*** that defines an external link to the root group of the corresponding file. This simple concept enables users to interact with the data as if it were located in a single file while the data is physically being stored distributed across many files.

In addition to the format-specific modules described so far, we use the generic ***AnnotationDataGroup*** (see Sec. 3.3.2) managed type for management and storage of collections of annotations associated with raw data and processed data (Sec. 3.3.2). We also use the generic ***ManagedObjectFile*** module (see Sec. 3.1.1) to support modular storage of managed objects in separate HDF5 files (which are in turn included in the parent via an external links). This allows users to flexibly store and share analytics as independent files while at the same time making the results easily accessible from the main data file and limiting the need for large-scale updates to the main file.

We will next discuss the storage of voltage recordings over time across multiple spatial distributed sensors via the ***BrainDataECoG*** and ***BrainDataECoGProcessed*** modules in more detail. For further details on the data organization we refer the interested reader to the specification documents of the data format shown in Supplement 2 (pp 35 – 62). Figure 1 also shows an abbreviated version of the specification document, listing all current managed object types in blue.

#### 4.1.2 Storing ECoG Data

A central application in neuroscience data is the acquisition and storage of voltage recordings over time across multiple spatial distributed sensors, e.g., via electrocorticography (ECoG), multi-channel electrophysiology from silicon shanks or Utah arrays. In the following we focus in particular on electrocorticography (ECoG) data collected from neurosurgical patients during speech production, however, we intend to extend these capabilities to other use-cases as well—such as physiology data collected in model species during standard sensory, motor, and cognitive neuroscience tasks—and the format has been designed with this extensibility in mind.

The ***BrainDataECoG*** module defines a managed group in HDF5 that serves as a container to collect all data pertaining to the voltage recordings in a single location. The primary dataset ***raw_data*** defines a two-dimensional, *space × time* array storing electrical recordings in units of *V olts*. Auxiliary information about the data, e.g, the ***sampling_rate***, in *Hz*, the ***unit*** of *V olts*, and the spatial ***layout*** of the electrodes are stored as additional datasets and attributes.

The ***raw_data*** is also further characterized via a series of dimensions scales describing: **1)** the identifier of electrodes (e.g. linear channel index from DAQ) (***electrode_id***), **2)** the sample time in milliseconds (***time_axis***), and optionally **3)** the anatomical name (***anatomy_name***) and integer id (***anatomy_id***) of the spatial region where each electrode is located. In addition, the ***BrainDataECoG*** API provides convenient functions to allow users to easily add custom dimension scales to the data. Dimension scales are described by: **1)** a data array with the scale’s data, **2)** the name of the unit of the data values, **3)** a human-readable description of the contents of the scale, **4)** the name of the scale, and **5)** the axis with which the scale is associated. The ability to easily generate custom dimension scales enables users to conveniently associate additional descriptions with the data, e.g., scales describing the classification of electrodes or time values into unique groups/clusters or to encode the occurrence of different events in time, such as, speech events or neural spikes among many others. Dedicated functions for look-up and retrieval of all or select dimensions scales—including all auxiliary data, e.g., the units or description of the scale(s)—ease the integration and use of dimensions scales for analytics.

Dimensions scales are limited in that they are one-dimensional in nature—specifically, even though the scale’s dataset may be an arbitrary n-dimensional array, the data is strictly associated with a particular dimension of the main dataset—and are not well-suited to describe complex structures, such as, multi-level data classifications with overlapping clusters. We, therefore, use the ***/data/internal/ecog_data_#/annotations_# : AnnotationDataGroup*** module (see Sec. 3.3.2) for storage and management of complex data annotations. The anatomical data, e.g., is automatically stored both via a dimension scale as well as annotations to facilitate the use of the anatomy in advanced analytics. The ***BrainDataECoG*** API also provides a number of convenience functions to assist with the interaction with and creation of custom collections of data annotations for the ***raw_data***. Annotations play a critical role in advanced analytics based on the classification of the data, e.g., based on the occurrence of events in time such as neural spikes or speech events. The definition of speech events in particular depends heavily on the ability to define many different types of annotations in conjunction with complex user-defined metadata associated with the annotations. For example, speech events occur at a broad range of nested classes, ranging from individual phonemes to syllables, words, and sentences etc.. The same speech event can occur arbitrary often during the course of an experiment—e.g, patient says ‘baa’—and different events can overlap—e.g., the sound ‘baa’ is part of the words ‘bad’. The ability to query annotations to locate particular speech events and subsequently analyze the data with such events is critical to the study of neural activity during speech production. Data annotation provides an ideal framework for storage and analysis of many derived classifications of electrical recordings, for example to define the occurrence of neural spikes via spike sorting.

As described earlier, the creation of managed objects is standardized, i.e., to create a new ***BrainDataECoG*** managed object we simply call the ***BrainDataECoG.create(…)*** function. All required data structures are initialized during the creation process, ensuring that the data file is always valid. Other, optional structures (e.g, the anatomy) may be saved directly during the creation or added later. To ease the use of ***BrainDataECoG*** during data acquisition, the create process allows the raw data and associated dimension scales to be initialized as empty datasets. As new recordings are acquired over time the ***raw_data*** and associated dimensions scales are then automatically expanded to accommodate the new data. With this so-called *auto-expand-data* feature enabled we can, for example do the following:

~~~
>>> from brain.dataformat.brainformat import BrainDataFile, BrainDataECoG
~~~

~~~
>>> import numpy
~~~

~~~
>>> brainfile = BrainDataFile.create(’testfile.h5’) # Create the file and initialize the data hierarchy
~~~

~~~
>>> internal_data = brainfile.data().internal() # Get the managed object for storing internal data
~~~

~~~
>>> ecog_data = BrainDataECoG.create(parent_object=internal_data, # Add to /data/internal
~~~

~~~
ecog_data_shape=(32,0), # Empty recording for 32 electrodes ecog_data_type=’f’, # Store float data values
~~~

~~~
chunks=True) # Store the data using chunking
~~~

~~~
>>> **ecog_data.set_auto_expand(True)** # Enable auto expansion
~~~

~~~
>>> **ecog_data[:, 0:1000] =** numpy.arange(32*1000).reshape(32,1000) # Add new data
~~~

Note when adding the new data, the shape of our ECoG dataset is automatically expanded to 32 *×* 1000 and all one-dimensional dimension-scales that are associated with the time axis are automatically expanded to match the new data shape so that we can also conveniently update the data of dimension scales without having to resize the datasets manually. As the above example illustrates, the ***BrainDataECoG*** API provides a convenient interface that allows us to directly interact with the primary ***raw_data*** via array slicing while auxiliary data, e.g., the sampling rate, layout, annotations etc., can be easily retrieved via corresponding access functions or key-based slicing(similar to Python dictionaries).

#### 4.1.3 Storing Processed ECoG Data

In practice, ECoG and other temporal voltage recordings across multiple sensors, are often further processed to extract specific, fixed-length tokens/features (e.g, phonemes or task trials) from the data. As a result the data is often reorganized as a three-dimensional array of *space × time × token*. The ***BrainDataECoGProcessed*** module is derived from ***BrainDataECoG*** and extends it to support storage of such processed data. Specifically: **1)** the primary dataset is extended by a third dimension to store the different *channels* and the dataset is renamed to ***processed_data***, **2)** a set of new optional dimension scales are specified to describe the *f requency*_*bands*, *token*_*id*, and *token*_*name*. Similar to the anatomy data, token data is stored both as dimension scales as well as via metadata-rich, searchable annotations to facilitate data analysis.

Figure 9 shows an example visualization of a processed ECoG dataset stored using our proposed data format. The tree view on the left shows the file structure, including all datasets associated with the ***/data/internal/ecog_data_processed_#*** group. The table view on the right then shows the contents of the primary ***processed_data*** dataset and the curve plot shows the voltage signal over time for a select set of tokens/electrodes. The properties view at the bottom then shows the shape, data type, and attributes associated with the main dataset.

## 5 CONCLUSIONS AND FUTURE WORK

Neuroscience is facing an incredible big data challenge. Efficient and easy-to-use data standards are a critical foundation to solving this challenge by enabling efficient storage, management, sharing, and analysis of complex neuroscience data. Standardizing neuroscience data is as much about defining common schema and ontologies for organizing and communicating data as it is about defining basic storage layouts for specific data types. Arguably, the focus of a neuroscience-oriented data standard should be on addressing the application-centric needs of organizing scientific data and metadata, rather than on reinventing file storage and format methods. For the development of BRAINformat we have used HDF5 as the basic storage format, because it already satisfies a broad range of the more basic format requirements.

The complexity and variety of experiments and the diversity of data types and acquisition modalities used in neuroscience make the creation of a general, all-encompassing data standard a daunting—if not futile—task. We have introduced the concept of *Managed Objects* (and *Managed Types*), which—in combination with an easy-to-use, formal format-specification document standard and API—enables us to divide & conquer the data standardization problem in a modular and extensible fashion. Format components specified using these concepts can be easily reused and extended and the format-compliance of file objects can be easily verified using the the BRAINformat library. The format specification API and managed object API implemented in BRAINformat are not specific to neuroscience, but define application-independent design concepts that enable us to efficiently create application-oriented data format modules. Based on these concepts we have developed an extensible data standard for neuroscience data that is portable, scalable, extensible, self-describing, and that supports self-contained (single-file) and modular (multiple-linked-files) storage.

We have also introduced a novel data format module and API for storage and management of advanced data annotations, enabling scientists to further characterize and organize data subsets via additional metadata descriptions. Additionally, we described the novel concept of relationship attributes for modeling and use of structural and semantic relationships between primary storage objects—including advanced index map relationships based on the concept of relationship chains. Although these features are available through an API, the data stored in the format is fully specified and human readable, so that domain scientists can access the data even without our API. These advanced capabilities fill critical gaps in the portfolio of available tools for creating advanced data standards for modern scientific data. The BRAINformat library is open source, has detailed developer documentation and user tutorials, and is freely available at: https://bitbucket.org/oruebel/brainformat.

In our future work we plan to extend the BRAINformat via advanced support for metadata ontology and data type specification capabilities and efficient metadata search, as well as expansion of the data annotation modules by supporting additional data selection schema and representations. We will develop capabilities to enable linking and interaction with external data stored in third-party formats (e.g. movies or images) and will develop additional data modules needed to provide a broader coverage of use cases in neuroscience research.

For concreteness, so far we have focused application of BRAINformat to electrocorticography data collected from neurosurgical patients during speech production. At the surface, it may appear that this is a specialization that hinders the general applicability of our work to the broader neuroscience community. However, the electrocorticography data shares many requirements with standard electrophysiology data collected by the community: storage of voltage recordings over time across multiple spatial distributed sensors with heterogenous geometries, complex and multi-tiered task descriptions, post-hoc processing of raw data to extract the signal of interest, the association of physiology data with multi-modal data streams collected simultaneously by other devices, the linkage of data associated with the same ‘task’ across multiple sessions, and the necessity to store rich meta-data to make sense of it all. Use of our format as input to popular spike-sorting algorithms, such as KlustaKwik [7], should be straightforward. Importantly, the utilization of metadata-rich Annotations are a natural way to encode the occurrence of spikes and associated parameters in the context of the original data, while relationship attributes provide an ideal foundation for recording relationships between analytics and other data. Therefore, application of BRAINformat to physiology data collected in model species during standard sensory, motor, and cognitive neuroscience tasks should be straightforward. Indeed, this has been our goal all along.

## ACKNOWLEDGMENTS

This work was supported by Laboratory Directed Research and Development (LDRD) funding from Berkeley Lab, provided by the Director, Office of Science, of the U.S. Department of Energy under Contract No. DE-AC02-05CH11231. This research used resources of the National Energy Research Scientific Computing Center, a DOE Office of Science User Facility supported by the Office of Science of the U.S. Department of Energy under Contract No. DE-AC02-05CH11231. We would like to thank Fritz Sommer, Jeff Teeters, Annette Greiner for helpful discussions. We would like to thank the members of the Chang Lab (UCSF) and Denes Lab (LBNL) for helpful discussions, data, and support.

## LEGAL DISCLAIMER

This document was prepared as an account of work sponsored by the United States Government. While this document is believed to contain correct information, neither the United States Government nor any agency thereof, nor The Regents of the University of California, nor any of their employees, makes any warranty, express or implied, or assumes any legal responsibility for the accuracy, completeness, or usefulness of any information, apparatus, product, or process disclosed, or represents that its use would not infringe privately owned rights. Reference herein to any specific commercial product, process, or service by its trade name, trademark, manufacturer, or otherwise, does not necessarily constitute or imply its endorsement, recommendation, or favoring by the United States Government or any agency thereof, or The Regents of the University of California. The views and opinions of authors expressed herein do not necessarily state or reflect those of the United States Government or any agency thereof or The Regents of the University of California.

## Footnotes

1 HDF users – https://www.hdfgroup.org/HDF5/users5.html

2 NeXus Definition Language (NXDL) – http://download.nexusformat.org/doc/html/nxdl.html

3 Orca slides presented at NWB: http://crcns.org/files/data/nwb/h1/NWBh1_09_Keith_Godfrey.pdf

4 Neurodata without Borders – https://crcns.org/NWB

5 http://crcns.org/NWB/hackathon-1

